# MousiPLIER: A Mouse Pathway-Level Information Extractor Model

**DOI:** 10.1101/2023.07.31.551386

**Authors:** Shuo Zhang, Benjamin J. Heil, Weiguang Mao, Maria Chikina, Casey S. Greene, Elizabeth A. Heller

## Abstract

High throughput gene expression profiling is a powerful approach to generate hypotheses on the underlying causes of biological function and disease. Yet this approach is limited by its ability to infer underlying biological pathways and burden of testing tens of thousands of individual genes. Machine learning models that incorporate prior biological knowledge are necessary to extract meaningful pathways and generate rational hypothesis from the vast amount of gene expression data generated to date. We adopted an unsupervised machine learning method, Pathway-level information extractor (PLIER), to train the first mouse PLIER model on 190,111 mouse brain RNA-sequencing samples, the greatest amount of training data ever used by PLIER. mousiPLER converted gene expression data into a latent variables that align to known pathway or cell maker gene sets, substantially reducing data dimensionality and improving interpretability. To determine the utility of mousiPLIER, we applied it to a mouse brain aging study of microglia and astrocyte transcriptomic profiling. We found a specific set of latent variables that are significantly associated with aging, including one latent variable (LV41) corresponding to striatal signal. We next performed k-means clustering on the training data to identify studies that respond strongly to LV41, finding that the variable is relevant to striatum and aging across the scientific literature. Finally, we built a web server (http://mousiplier.greenelab.com/) for users to easily explore the learned latent variables. Taken together this study provides proof of concept that mousiPLIER can uncover meaningful biological processes in mouse transcriptomic studies.

**Significance statement:** Analysis of RNA-sequencing data commonly generates differential expression of individual genes across conditions. However, genes are regulated in complex networks, not as individual entities. Machine learning models that incorporate prior biological information are a powerful tool to analyze human gene expression. However, such models are lacking for mouse despite the vast number of mouse RNA-seq datasets. We trained a mouse pathway-level information extractor model (mousiPLIER). The model reduced the data dimensionality from over 10,000 genes to 196 latent variables that map to prior pathway and cell marker gene sets. We demonstrated the utility of mousiPLIER by applying it to mouse brain aging data and developed a web server to facilitate the use of the model by the scientific community.

## Introduction

Over the last decade, scientists have generated an astronomical amount of brain gene expression data (Carulli et al., 1998; Anders and Huber, 2010; Costa-Silva et al., 2017; Keil et al., 2018; Zhang et al., 2021). Differential gene expression analysis of high throughput RNA-sequencing data is commonly applied to interrogate the relative enrichment of a single transcript across samples. However, genes are regulated in complex networks, rather than as individual entities. Furthermore, gene expression profiling studies are limited in statistical power, as they tend to examine relatively few samples compared to the number of expressed transcripts and increasing the number of samples can be prohibitively expensive.

Machine learning models that incorporate prior pathway information have shown great power in analyzing human gene expression. To this end we apply an unsupervised learning method that (1) reduces the dimensionality and/or (2) incorporates additional published gene expression datasets. Unsupervised machine learning is a method that defines the structure of “unlabeled data”, for which information on the biological context and experimental conditions is removed. Such methods are well-suited for gene expression data, and are often used for tasks such as reducing the dimensionality of expression datasets (Hotelling, 1933; der Maaten and Hinton, 2008; McInnes et al., 2018), clustering samples (Oyelade et al., 2016; Chen et al., 2020), or learning shared expression patterns across experiments (Tan et al., 2016; Handl et al., 2019). While unsupervised machine learning models are capable of analyzing large amounts of unlabeled expression data, many of them don’t explicitly encode prior biological knowledge to encourage the model to learn biologically meaningful patterns of gene expression over technical ones.

A novel approach, the modeling framework pathway-level information extractor (PLIER) (Mao et al., 2019), is built explicitly to work on expression data, and uses matrix factorization to incorporate prior biological knowledge in the form of sets of genes corresponding to biological pathways or cell type markers. This approach converts gene expression data into a series of values called “latent variables” (LVs) that correspond to potentially biologically relevant combinations of differentially expressed genes. PLIER learns diverse biological pathways from entire compendia of expression data and can transfer that knowledge to smaller studies, such as MultiPLIER (Taroni et al., 2019). However, PLIER models are largely trained on a single dataset rather than a compendium (Rubenstein et al., 2020; Stogsdill et al., 2022; Zhang et al., 2022), and past MultiPLIER runs have only trained models with up to tens of thousands of samples (Taroni et al., 2019; Banerjee et al., 2020).

To expand the application and utility of PLIER for identifying meaningful biological pathways from gene expression data, we trained a PLIER model on a compendium of mouse gene expression data. In doing so we trained the first mouse compendium PLIER model (mousiPLIER), on the greatest amount of training data (190,111 samples) ever used by this model. We demonstrated successful optimization of the model training, which generated hypotheses on regulation of mouse brain aging. A further innovation applied k-means clustering in the latent variable space to identify the microglia-associated latent variables that corresponded to aging-related changes in the training data. Finally, to maximize widespread usability of mousiPLIER, we built a web server that allows others to visualize the results and find patterns in the data based on their own latent variables of interest. Going forward, this model and its associated web server will be a useful tool for better understanding mouse gene expression.

## Materials and Methods

### Data

We began by downloading all the mouse gene expression data in Recount3, along with its corresponding metadata (Wilks et al., 2021). We then removed the single-cell RNA-seq data from the dataset to ensure our data sources were consistent across samples and studies. Next, we filtered the expression data, keeping only genes that overlapped between Recount3 and our prior knowledge gene sets. Finally, we RPKM-transformed the expression using gene lengths from the Ensembl BioMart database (Howe et al., 2021) and Z-scored the expression data to ensure a consistent range for the downstream PLIER model.

For our prior-knowledge gene sets we used cell type marker genes from CellMarker (Zhang et al., 2019), pathway gene sets from Reactome (Gillespie et al., 2022), and manually curated brain marker genes from Allen Mouse Brain Atlas (https://mouse.brain-map.org) (Lein et al., 2007). We selected cell type marker genes corresponding to all available mouse cell types within the CellMarker database. For mouse biological pathways, we downloaded pathway information from the Reactome database. More specifically, we processed the files “Ensembl2Reactome_All_ Levels.txt”, “ReactomePathways.txt”, and “ReactomePathwaysRelation.txt”, selecting only pathways using mouse genes, filtering out all pathways with fewer than 5 genes present, and keeping only pathways that were leaf nodes on the pathway hierarchy. Because we were interested in mouse brains in particular, we rounded out our set of prior information by manually selecting marker genes for the striatum, midbrain, and cerebellum. In total, we used 1,003 prior knowledge pathways when training our model.

### PLIER

We began the PLIER pipeline by precomputing the initialization for PLIER with incremental PCA in scikit-learn (Pedregosa et al., 2011). We then used the expression compendium, prior knowledge gene sets, and PCA initializations to train a PLIER model. The resulting task took two days to run and yielded 196 latent variables.

### RNA-seq processing

RNA-seq reads from mouse microglia and astrocytes (Pan et al., 2020) were mapped to mm10 reference genome using STAR (v2.7.1a)(Dobin et al., 2013) with parameters: --outFilterMismatchNmax 3 --outFilterMultimapNmax 1 --alignSJoverhangMin 8. Gene level read counts were prepared using featureCounts (subread v1.6.1)(Liao et al., 2014). The gene annotation file used in featureCounts was download from recount3 (https://rna.recount.bio/docs/raw-files.html#annotation-files).

### LV significance for mouse aging RNA-seq data

We first transformed the mouse aging expression data from gene space to latent variable space using a custom Python script. To determine which latent variables were associated with experimental conditions, we used a linear model by treating development stages (in month) as numerical variables. To correct the p-values for multiple testing, we used the Benjamini-Hochberg procedure (FDR)(Benjamini and Hochberg, 1995). LVs with FDR < 0.05 were considered to be significantly associated.

### Clustering

We selected the latent variables significantly associated with aging in mouse microglia as a biological starting point. We then used these latent variables to query the training data and see which studies seemed associated with the same biological signals. To do so, we used *k*-means clustering with a *k* of 2, to look for experiments where there was some experimental condition that affected the latent variable. We then ranked the top ten studies based on their silhouette scores, and looked to see which conditions were associated with relevant experimental variables.

### Hardware

The PLIER model training was performed on the Penn Medicine high performance computing cluster. The full pipeline takes around two weeks to run, with the main bottlenecks being the Recount3 data download, which takes one week to run, and training the PLIER model, which takes two days on a compute node with 250GB of RAM.

### Web Server

The web server for visualizing the results was built on top of the ADAGE web app framework (Tan et al., 2017). The main changes we made were to substitute the latent variables and gene sets from our trained PLIER model and to forgo uploading the input expression data as the mouse compendium we used was much larger than the input expression for ADAGE.

### Data and code availability

All data and code used in this study can be found at https://github.com/greenelab/mousiplier.

## Results

### MousiPLIER learned latent variables with ideal pathway-level and gene-level sparsity

We trained mousiPLIER using on-disk PCA implementation to initialize PLIER, modified the pipeline to work with mouse data, and used a high-memory compute node to manage the size of the matrix decomposition (see Materials and Methods). The resulting model had 196 latent variables with ideal pathway-level and gene-level sparsity. The per-latent variable distribution had an average of 65% sparsity, such that the latent variables tended to use only around 35% of the genes in the training data (Fig. 1A). While many of the latent variables corresponded to no pathways, indicating signals in the training data not passed in as prior knowledge, those that remained corresponded to few pathways (Fig. 1B). This optimal pathway-level and gene-level sparsity allowed us to interrogate individual latent variables that corresponded to a small number of biological functions.

**Figure 1.**
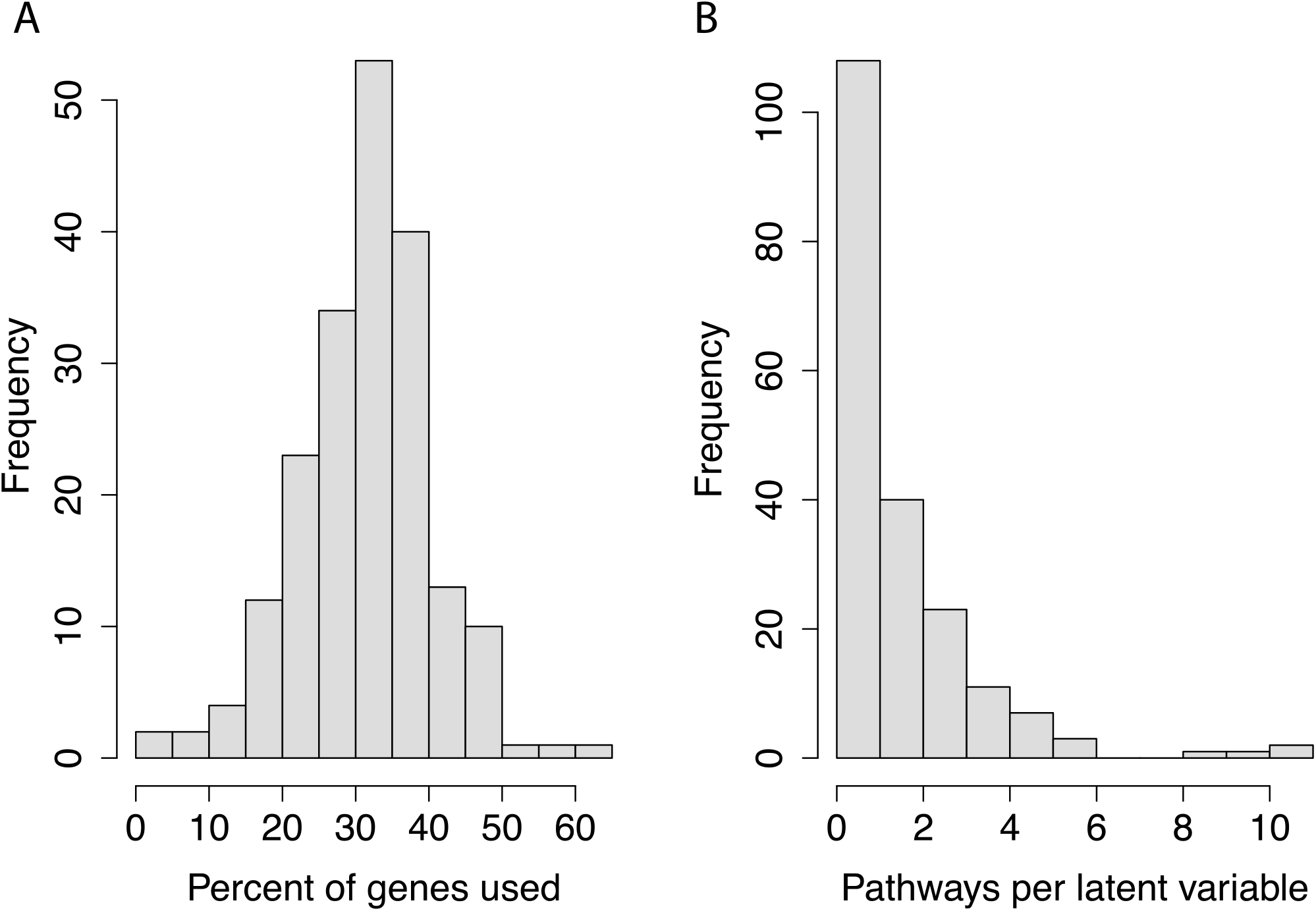
mousiPLIER learned latent variables with ideal pathway-level and gene-level sparsity. (A) The distribution of the percentage of genes from the training set used per latent variable. (B) The distribution of the number of prior knowledge gene sets used per latent variable.

### MousiPLIER identified LVs associated with aging

We next interrogated brain-relevant latent variables that our mousiPLIER learned from the compendium. We analyzed an individual study on mouse brain aging (Pan et al., 2020). This study measures wild-type and Alzheimer’s disease (APP-PS1) mouse gene expression in microglia and astrocytes at five ages across adulthood. We first projected the RNA-seq data from gene space to mousiPLIER LV space. Then, we used a linear model to identify the latent variables that changed significantly across developmental aging. We found that mousiPLIER identified a specific set of LVs in each condition in the study (Fig. 2A). To narrow down the scope of the analysis, we next validated the biological relevance of the latent variables associated with wild-type microglial cells.

**Figure 2.**
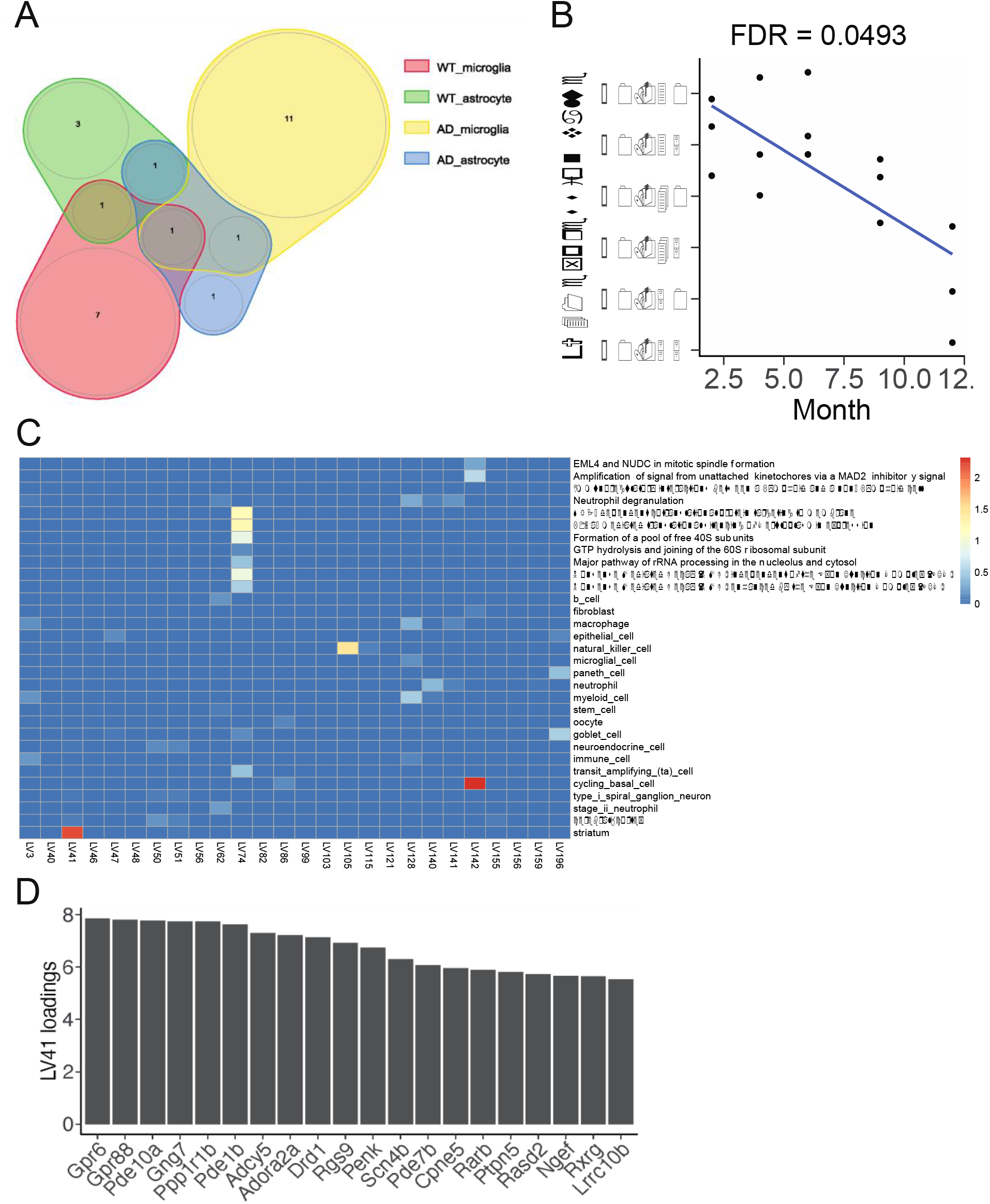
MousiPLIER identified LVs associated with aging. (A) Venn diagram showing the number of significant LVs and their overlap across cell types and experimental conditions. (B) LV41 is deceased significantly in wild-type microglia during aging. (C) A heatmap showing significant LV-associated biological pathways or cell type markers. A linear model is used to test the effect of aging on each LV. P-values are adjusted for multiple comparisons using Benjamini-Hochberg method. A LV is differentially expressed if FDR < 0.05. LV: latent variable; AD, Alzheimer’s disease; WT, wild-type.

### Latent variable 41 demonstrated the biological relevance of mousiPLIER latent variables

Having identified microglia-associated latent variables of interest, we next sought to validate the relevance of this gene set by finding which studies in the mousiPLIER training data responded strongly to them. To do so, we developed a novel method to rank studies based on their latent variable weights. More precisely, we performed *k*-means clustering with a *k* of two on each study in each latent variable space, and ranked studies by their silhouette scores, a metric measuring the degree to which clusters are separated from each other. Using this approach, we identified studies that contained samples distinguishable by their values for our latent variables of interest.

We focused this approach on latent variable 41, which decreased throughout aging in wild-type microglia and contained genes functionally associated with striatal cell type specificity (Fig. 2B-D). We found that many of the studies with the highest silhouette scores for latent variable 41 indicated processes occurring in the brain (Fig. 3A). We dug deeper into which specific samples were present in each cluster and found that latent variable 41 was in fact learning something brain- (and more specifically striatum-) related (Fig. 3).

**Figure 3.**
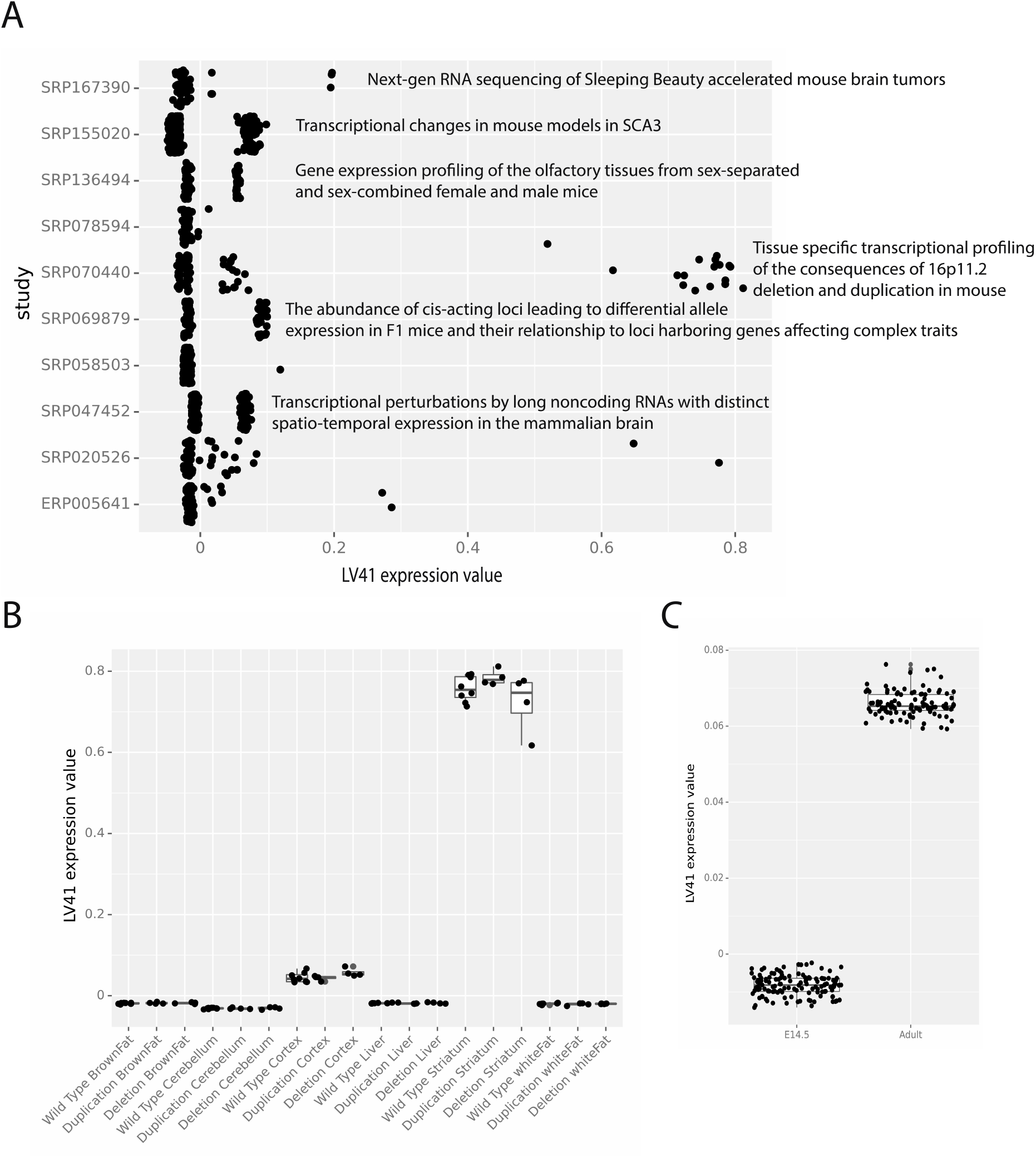
Latent variable 41 demonstrated the biological relevance of mousiPLIER latent variables. (A) Studies with the ten highest silhouette scores after clustering according to LV41 expression values. (B) LV41 expression values are higher for striatal tissue than other tissues in SRP070440. (C) Effects of development on LV41 (D) Top 20 genes with highest weight associated with LV41. LV: latent variable.

For example, in a study to delineate tissue-specific transcriptional consequences of copy number variant within 16p11.2, a common cause of autism spectrum disorder, gene expression data were profiled from three genotypes (wild type, deletion, and duplication of the 16p11.2 region) and six tissues (brown fat, liver, white fat, cerebellum, cortex, and striatum) (https://www.ncbi.nlm.nih.gov/geo/query/acc.cgi?acc=GSE76872). The LV experimental values in the striatal samples, irrespective of the genotype, clearly stand apart from the other tissues (Fig. 3B). Additionally, in a study to investigate transcriptional effects of selected long noncoding RNAs (lncRNAs), mRNA expression is generated from embryonic and adult whole brains of wild-type and lncRNA knockout mouse (Goff et al., 2015). The two groups distinguished by latent variable 41 consist of brain samples from embryonic and adult mice (Fig. 3C), supporting the association between latent variable 41 and aging found in the study (Pan et al., 2020) we used to derive the latent variables.

### Web Server

To allow others to independently examine mousiPLIER learned latent variables, we developed a web server at http://mousiplier.greenelab.com/. This server allows users to list the genes present in, visualize which experiments had high cluster scores for, and see which biological pathways participate in each latent variable (Fig. 4).

**Figure 4.**
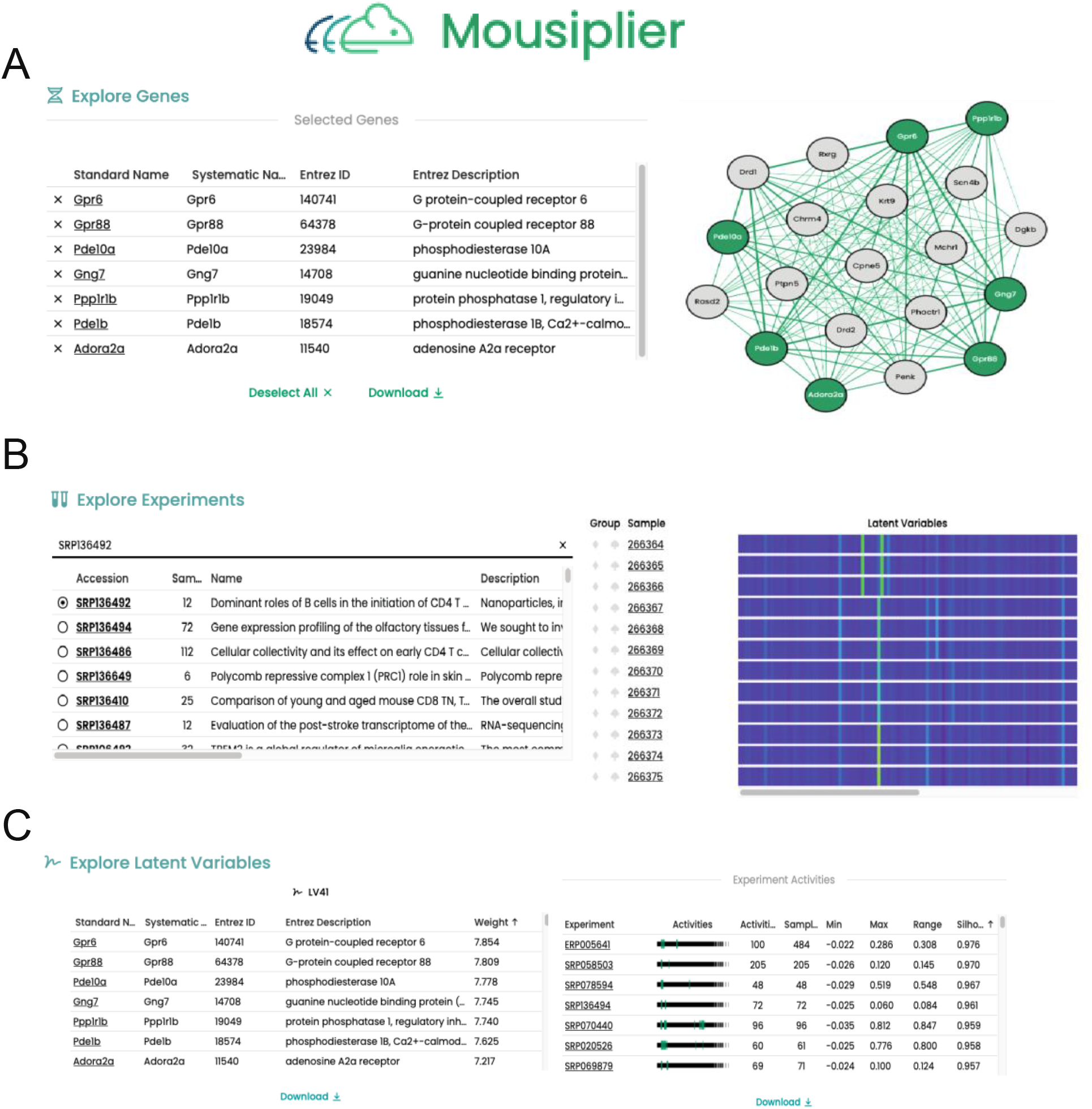
Snapshot of mousiPLIER web server showing its functions. (A) An example to visualize the gene network of LV41 associated genes. (B) An example to explore experiments and view samples’ activities in mousiPLIER latent space. (C) An example to explore latent variables. Gene weights and the clustering of experiments for each LV is displayed and can be easily downloaded.

## Discussion/Conclusion

In this paper we developed mousiPLIER and established proof of concept for training extremely large PLIER models on mouse data. The learned latent variables mapped to various biological processes and cell types. Further, we applied a novel approach for surfacing latent-variable relevant experiments from an expression compendium. Specifically, we clustered training experiments based on latent variable values, allowing us to query a large compendium for experiments pertaining to mouse striatal aging. Finally, we created a web server to make the model’s results more easily accessible to other scientists.

While we successfully transformed a study from outside the training data into the latent space and identified study-specific latent variables, application of mousiPLIER was not universally successful across transcriptomic studies. This may be due to lack of similar samples in training compendium, too few samples in the study of interest to generate sufficient statistical power, or other factors. In these cases, there isn’t yet a method to select meaningful latent variables to guide downstream analyses.

Finally, as a linear model PLIER can only approximate non-linear relationships between the genes used to train the model and the learned biological pathways. While we do not expect this to have a large impact (Heil et al., 2022), incorporating prior knowledge into non-linear models such as neural networks is an exciting field of research and a potential improvement for the MultiPLIER and mousiPLIER frameworks. Going forward, our model and web server will allow scientists to explore the latent space of their own experiments and learn about relevant biological pathways and cell types.

## References

Anders S, Huber W (2010) Differential expression analysis for sequence count data. Nat Preced:1.

Banerjee J, Allaway RJ, Taroni JN, Baker A, Zhang X, Moon CI, Pratilas CA, Blakeley JO, Guinney J, Hirbe A, others (2020) Integrative analysis identifies candidate tumor microenvironment and intracellular signaling pathways that define tumor heterogeneity in NF1. Genes (Basel) 11:226.

Benjamini Y, Hochberg Y (1995) Controlling the false discovery rate: a practical and powerful approach to multiple testing. J R Stat Soc B 57:289–300 Available at: http://www.stat.purdue.edu/~doerge/BIOINFORM.D/FALL06/Benjamini and Y FDR.pdf%5Cn http://engr.case.edu/ray_soumya/mlrg/controlling_fdr_benjamini95.pdf.

Carulli JP, Artinger M, Swain PM, Root CD, Chee L, Tulig C, Guerin J, Osborne M, Stein G, Lian J, others (1998) High throughput analysis of differential gene expression. J Cell Biochem 72:286–296.

Chen R, Yang L, Goodison S, Sun Y (2020) Deep-learning approach to identifying cancer subtypes using high-dimensional genomic data. Bioinformatics 36:1476–1483.

Costa-Silva J, Domingues D, Lopes FM (2017) RNA-Seq differential expression analysis: An extended review and a software tool. PLoS One 12:e0190152.

der Maaten L, Hinton G (2008) Visualizing data using t-SNE. J Mach Learn Res 9.

Dobin A, Davis CA, Schlesinger F, Drenkow J, Zaleski C, Jha S, Batut P, Chaisson M, Gingeras TR (2013) STAR: Ultrafast universal RNA-seq aligner. Bioinformatics 29:15–21.

Gillespie M, Jassal B, Stephan R, Milacic M, Rothfels K, Senff-Ribeiro A, Griss J, Sevilla C, Matthews L, Gong C, others (2022) The reactome pathway knowledgebase 2022. Nucleic Acids Res 50:D687--D692.

Goff LA, Groff AF, Sauvageau M, Trayes-Gibson Z, Sanchez-Gomez DB, Morse M, Martin RD, Elcavage LE, Liapis SC, Gonzalez-Celeiro M, others (2015) Spatiotemporal expression and transcriptional perturbations by long noncoding RNAs in the mouse brain. Proc Natl Acad Sci 112:6855–6862.

Handl L, Jalali A, Scherer M, Eggeling R, Pfeifer N (2019) Weighted elastic net for unsupervised domain adaptation with application to age prediction from DNA methylation data. Bioinformatics 35:i154.-i163.

Heil BJ, Crawford J, Greene CS (2022) The Effects of Nonlinear Signal on Expression-Based Prediction Performance. bioRxiv.

Hotelling H (1933) Analysis of a complex of statistical variables into principal components. J Educ Psychol 24:417.

Howe KL, Achuthan P, Allen J, Allen J, Alvarez-Jarreta J, Amode MR, Armean IM, Azov AG, Bennett R, Bhai J, others (2021) Ensembl 2021. Nucleic Acids Res 49:D884--D891.

Keil JM, Qalieh A, Kwan KY (2018) Brain transcriptome databases: a user’s guide. J Neurosci 38:2399–2412.

Lein ES, Hawrylycz MJ, Ao N, Ayres M, Bensinger A, Bernard A, Boe AF, Boguski MS, Brockway KS, Byrnes EJ, others (2007) Genome-wide atlas of gene expression in the adult mouse brain. Nature 445:168–176.

Liao Y, Smyth GK, Shi W (2014) featureCounts: an efficient general purpose program for assigning sequence reads to genomic features. Bioinformatics 30:923–930.

Mao W, Zaslavsky E, Hartmann BM, Sealfon SC, Chikina M (2019) Pathway-level information extractor (PLIER) for gene expression data. Nat Methods 16:607–610.

McInnes L, Healy J, Melville J (2018) Umap: Uniform manifold approximation and projection for dimension reduction. arXiv Prepr arXiv180203426.

Oyelade J, Isewon I, Oladipupo F, Aromolaran O, Uwoghiren E, Ameh F, Achas M, Adebiyi E (2016) Clustering algorithms: their application to gene expression data. Bioinform Biol Insights 10:BBI--S38316.

Pan J, Ma N, Yu B, Zhang W, Wan J (2020) Transcriptomic profiling of microglia and astrocytes throughout aging. J Neuroinflammation 17:1–19.

Pedregosa F, Varoquaux G, Gramfort A, Michel V, Thirion B, Grisel O, Blondel M, Prettenhofer P, Weiss R, Dubourg V, Vanderplas J, Passos A, Cournapeau D, Brucher M, Perrot M, Duchesnay E (2011) Scikit-learn: Machine Learning in Python. J Mach Learn Res 12:2825–2830.

Rubenstein AB, Smith GR, Raue U, Begue G, Minchev K, Ruf-Zamojski F, Nair VD, Wang X, Zhou L, Zaslavsky E, others (2020) Single-cell transcriptional profiles in human skeletal muscle. Sci Rep 10:1–15.

Stogsdill JA, Kim K, Binan L, Farhi SL, Levin JZ, Arlotta P (2022) Pyramidal neuron subtype diversity governs microglia states in the neocortex. Nature 608:750–756.

Tan J, Hammond JH, Hogan DA, Greene CS (2016) ADAGE-Based Integration of Publicly Available Pseudomonas aeruginosa Gene Expression Data with Denoising Autoencoders Illuminates Microbe-Host Interactions. mSystems 1:e00025–15.

Tan J, Huyck M, Hu D, Zelaya RA, Hogan DA, Greene CS (2017) ADAGE signature analysis: differential expression analysis with data-defined gene sets. BMC Bioinformatics 18:1–15.

Taroni JN, Grayson PC, Hu Q, Eddy S, Kretzler M, Merkel PA, Greene CS (2019) MultiPLIER: a transfer learning framework for transcriptomics reveals systemic features of rare disease. Cell Syst 8:380–394.

Wilks C, Zheng SC, Chen FY, Charles R, Solomon B, Ling JP, Imada EL, Zhang D, Joseph L, Leek JT, others (2021) recount3: summaries and queries for large-scale RNA-seq expression and splicing. Genome Biol 22:1–40.

Zhang X, Lan Y, Xu J, Quan F, Zhao E, Deng C, Luo T, Xu L, Liao G, Yan M, others (2019) CellMarker: a manually curated resource of cell markers in human and mouse. Nucleic Acids Res 47:D721--D728.

Zhang Y, Thompson KN, Huttenhower C, Franzosa EA (2021) Statistical approaches for differential expression analysis in metatranscriptomics. Bioinformatics 37:i34.–i41.

Zhang Z, Zamojski M, Smith GR, Willis TL, Yianni V, Mendelev N, Pincas H, Seenarine N, Amper MAS, Vasoya M, others (2022) Single nucleus transcriptome and chromatin accessibility of postmortem human pituitaries reveal diverse stem cell regulatory mechanisms. Cell Rep 38:110467.

